# Investigating the impact of reference assembly choice on genomic analyses in a cattle breed

**DOI:** 10.1101/2021.01.15.426838

**Authors:** Audald Lloret-Villas, Meenu Bhati, Naveen Kumar Kadri, Ruedi Fries, Hubert Pausch

## Abstract

**Background:** Reference-guided read alignment and variant genotyping are prone to reference allele bias, particularly for samples that are greatly divergent from the reference genome. A Hereford-based assembly is the widely accepted bovine reference genome. Haplotype-resolved genomes that exceed the current bovine reference genome in quality and continuity have been assembled for different breeds of cattle. Using whole genome sequencing data of 161 Brown Swiss cattle, we compared the accuracy of read mapping and sequence variant genotyping as well as downstream genomic analyses between the bovine reference genome (ARS-UCD1.2) and a highly continuous Angus-based assembly (UOA_Angus_1).

**Results:** Read mapping accuracy did not differ notably between the ARS-UCD1.2 and UOA_Angus_1 assemblies. We discovered 22,744,517 and 22,559,675 high-quality variants from ARS-UCD1.2 and UOA_Angus_1, respectively. The concordance between sequence- and array-called genotypes was high and the number of variants deviating from Hardy-Weinberg proportions was low at segregating sites for both assemblies. More artefactual INDELs were genotyped from UOA_Angus_1 than ARS-UCD1.2 alignments. Using the composite likelihood ratio test, we detected 40 and 33 signatures of selection from ARS-UCD1.2 and UOA_Angus_1, respectively, but the overlap between both assemblies was low. Using the 161 sequenced Brown Swiss cattle as a reference panel, we imputed sequence variant genotypes into a mapping cohort of 30,499 cattle that had microarray-derived genotypes. The accuracy of imputation (Beagle R^2^) was very high (0.87) for both assemblies. Genome-wide association studies between imputed sequence variant genotypes and six dairy traits as well as stature produced almost identical results from both assemblies.

**Conclusions:** The ARS-UCD1.2 and UOA_Angus_1 assemblies are suitable for reference-guided genome analyses in Brown Swiss cattle. Although differences in read mapping and genotyping accuracy between both assemblies are negligible, the choice of the reference genome has a large impact on detecting signatures of selection using the composite likelihood ratio test. We developed a workflow that can be adapted and reused to compare the impact of reference genomes on genome analyses in various breeds, populations and species.

## Background

Representative reference genomes are paramount for genome research. A reference genome is an assembly of digital nucleotides that are representative of a species’ genetic constitution. Like the coordinate system of a two-dimensional map, the coordinates of the reference genome unambiguously point to nucleotides and annotated genomic features. Because the physical position and alleles of sequence variants are determined according to reference coordinates, the adoption of a universal reference genome is required to compare findings across studies. Otherwise, the conversion of genomic coordinates between assemblies is necessary [1]. Updates and amendments to the reference genome change the coordinate system.

Reference genomes of important farm animal species including cattle, pig and chicken were assembled more than a decade ago using bacterial artificial chromosome and whole-genome shotgun sequencing [2, 3, 4]. The initial reference genome of domestic cattle (Bos taurus taurus) was generated from a DNA sample of the inbred Hereford cow ≪L1 Dominette 01449≫ [3, 5]. An annotated bovine reference genome enabled systematic assessment and characterization of sequence variation within and between cattle populations using reference-guided alignment and variant detection [3, 6]. A typical genome-wide alignment of DNA sequences from a *B. taurus taurus* individual differs at between 6 and 8 million single nucleotide polymorphisms (SNPs) and small (*<* 50 bp) insertions and deletions (INDELs) from the reference genome [7, 8]. More variants are detected in cattle with greater genetic distance from the Hereford breed [9]. The bovine reference genome neither contains allelic variation nor nucleotides that are private to animals other than ≪L1 Dominette 01449≫. As a result, read alignments may be erroneous particularly at genomic regions that differ substantially between the sequenced individual and the reference genome [10]. The use of consensus reference genomes or variation-aware reference graphs may mitigate this type of bias [11, 12, 13].

The quality of reference genomes improved spectacularly over the past 15 years. Decreasing error rates and increasing outputs of long-read (*>* 10 Kb) sequencing technologies such as PacBio single molecule real-time (SMRT) [14] and Oxford Nanopore sequencing [15] revolutionised the assembly of reference genomes. Sophisticated genome assembly methods enable to assemble gigabase-sized and highly-repetitive genomes from long sequencing reads at high continuity and accuracy [16, 17, 18]. The application of “trio-binning” [19] facilitates the *de novo* assembly of haplotype-resolved genomes that exceed in quality and continuity all previously assembled reference genomes. This approach now offers an opportunity to obtain reference-quality genome assemblies and identify hitherto undetected variants in non-reference sequences, thus making the full spectrum of sequence variation amenable to genetic analyses [17, 19].

Reference-quality assemblies are available for Hereford (ARS-UCD1.2) [20], Angus (UOA_Angus_1) [17] and Highland cattle [21]. In addition, reference-quality assemblies are available for yak (*Bos grunniens*) [21] and Brahman (*Bos taurus indicus*) [17] which are closely related to taurine cattle. Any of these resources may serve as a reference for reference-guided sequence read alignment, variant detection and annotation. Linear mapping and sequence variant genotyping accuracy may be affected by the choice of the reference genome and the divergence of the DNA sample from the reference genome [22, 23, 24, 25]. It remains an intriguing question, which reference genome enables optimum read mapping and variant detection accuracy for a particular animal [11, 12, 13].

Here, we assessed the accuracy of reference-guided read mapping and sequence variant detection in 161 Brown Swiss (BSW) cattle using two highly continuous bovine genome assemblies that were created from Hereford (ARS-UCD1.2) and Angus (UOA_Angus_1) cattle. Moreover, we detect signatures of selection and perform sequence-based association studies to investigate the impact of the reference genome on downstream genomic analyses.

## Results

Short paired-end whole-genome sequencing reads of 161 BSW cattle (113 males, 48 females) were considered for our analysis. All raw sequencing data are publicly available at the Sequencing Read Archive of the NCBI [26] or the European Nucleotide Archive of the EMBL-EBI [27]. Accession numbers are listed in the Supplementary File 1: Table S1.

### Alignment quality and depth of coverage

Following the removal of adapter sequences, and reads and bases of low sequencing quality, between 173 and 1,411 million reads per sample (mean: 360 ± 165 million reads) were aligned to expanded versions of the Hereford-based ARS-UCD1.2 and the Angus-based UOA_Angus_1 assemblies that included sex chromosomal sequences and unplaced scaffolds (see Material and Methods) using a reference-guided alignment approach. The Hereford assembly is a primary assembly because it was created from a purebred animal [20]. The Angus assembly is haplotype-resolved because it was created from an Angus x Brahman cross using “trio-binning” [17]. The average number of reads per sample that aligned to sex chromosomes, the mitochondrial genome and unplaced contigs were slightly higher for UOA_Angus_1 (66 ± 39 million) than ARS-UCD1.2 (64 ± 38 million).

We considered the 29 autosomes to investigate alignment quality. The total length of the autosomes was 2,489,385,779 bp for ARS-UCD1.2 and 2,468,157,877 bp for UOA_Angus_1. An average number of 295 ± 131 and 293 ± 130 million reads per sample aligned to autosomal sequences of ARS-UCD1.2 and UOA_Angus_1, respectively. The slightly higher number of reads that mapped to ARS-UCD1.2 is likely due to its longer autosomal sequence. In order to ensure consistency across all analyses performed, we retained 263 ± 118 (89.28%) and 261 ± 117 (89.17%) uniquely mapped and properly paired reads (*i.e.*, all reads except those with a SAM-flag value of 1796) that had mapping quality higher than 10 (high-quality reads hereafter) per sample, as such reads qualify for sequence variant genotyping using the best practice guidelines of the Genome Analysis Toolkit (GATK) [28, 29] (Table 1). The number of reads that mapped to the autosomes but were discarded due to low mapping quality (either SAM-flag 1796 or MQ *<* 10) were almost identical (32 ± 20 million) for both assemblies (Supplementary File 2: Table S2). Most of the discarded reads (83.37% for ARS-UCD1.2 and 82.29% for UOA_Angus_1) were flagged as duplicates.

**Table 1.** Mapping statistics for the 161 BSW samples. Summary statistics extracted from the BAM files after aligning the samples to either the ARS-UCD1.2 or UOA_Angus_1 assembly. Uniquely mapped and properly paired reads with MQ > 10 are considered as high-quality reads. The percentage of autosomal reads that are high-quality reads is calculated per sample and per chromosome. Coverage of high-quality reads is calculated per sample and per chromosome.

| Parameter | Unit | ARS-UCD1.2 | UOA_Angus.1 |
| --- | --- | --- | --- |
| Autosomal reads | Million | 47,502 | 47,128 |
| | Million / sample | $295 \pm 131$ | $293 \pm 130$ |
| Autosomal high-quality reads | Million | 42,418 | 42,029 |
| | Million / sample | $263 \pm 118$ | $261 \pm 117$ |
| | % / sample | $89.28 \pm 5.06$ | $89.17 \pm 5.06$ |
| | % / chromosome | $89.28 \pm 0.34$ | $89.17 \pm 0.56$ |
| Coverage | fold / sample | $14.13 \pm 7.26$ | $14.11 \pm 7.25$ |
| | fold / chromosome | $14.13 \pm 0.14$ | $14.11 \pm 0.15$ |

The mean percentage of high-quality reads was slightly higher (0.10 ± 0.63) for the ARS-UCD1.2 than UOA_Angus_1 autosomes but greater differences existed at some chromosomes. The proportion of high-quality reads was higher for the ARS-UCD1.2 assembly than the UOA_Angus_1 assembly at 16 out of the 29 autosomes. The greatest difference was observed for chromosome 20, for which the proportion of high-quality reads was 2.03 percent points greater for the ARS-UCD1.2 assembly than the UOA_Angus_1 assembly (P = 4.5 × 10^−4^). Of 8.59 ± 3.81 and 8.69 ± 3.88 million reads that aligned to chromosome 20 of ARS-UCD1.2 and UOA_Angus_1, respectively, 7.66 ± 3.42 and 7.57 ± 3.38 million were high-quality reads. Among the 13 autosomes for which the percentage of high-quality reads was greater for the UOA_Angus_1 than ARS-UCD1.2 assembly, the greatest difference (0.75 percent points) was observed for chromosome 13.

Average genome coverage ranged from 8.8- to 62.4-fold per sample for both assemblies. The mean coverage of the BAM files was nearly identical for the ARS-UCD1.2 (14.13 ± 7.26) and UOA_Angus_1 (14.11 ± 7.25) assembly. Chromosome wise, no differences were detected (P = 0.36) across the two assemblies considered. The mean coverage was between 13.76 (chromosome 19) and 14.45 (chromosome 27) for ARS-UCD1.2 and between 13.76 (chromosome 19) and 14.52 (chromosome 14) for UOA_Angus_1.

### Sequence variant genotyping and variant statistics

Single nucleotide polymorphisms (SNPs), insertions and deletions (INDELs) were discovered from the BAM files following the GATK best practice guidelines [28, 29]. Using the HaplotypeCaller and GenotypeGVCFs modules of GATK, we detected 24,760,861 and 24,557,291 autosomal variants from the ARS-UCD1.2 and UOA_Angus_1 alignments, respectively, of which 22,744,517 (91.86%) and 22,559,675 (91.87%) high-quality variants were retained after applying site-level hard filtration using the VariantFiltration module of GATK (Supplementary File 3: Table S3). The mean transition/transversion ratio was 2.15 for the high-quality variants detected from either of the assemblies.

For 32.40 and 33.80% of the high-quality variants, the genotype of at least one out of 161 BSW samples was missing using the ARS-UCD1.2 and UOA_Angus_1 alignments, respectively. Across all chromosomes, the number of missing genotypes was slightly higher (P = 0.087) for variants called from UOA_Angus_1 than ARS-UCD1.2 alignments. The percentage of variants with missing genotypes was highest on chromosome 12 in both assemblies. At least one missing genotype was observed for 49.79 and 37.39% of the chromosome 12 variants for the UOA_Angus_1 and ARS-UCD1.2-called genotypes. Beagle [30] (version 4.1) phasing and imputation was applied to improve the genotype calls from GATK and impute the missing genotypes.

112 sequenced animals that had an average fold sequencing coverage of 13.47 ± 6.45 and 13.46 ± 6.44 when aligned to ARS-UCD1.2 and UOA_Angus_1, respectively, also had Illumina BovineHD array-called genotypes at 530,372 autosomal SNPs. We considered the microarray-called genotypes as a truth set to calculate non-reference sensitivity, non-reference discrepancy and the concordance between array-called and sequence-called genotypes (Table 2). The average concordance between array- and sequence-called genotypes was greater than 98 and 99.5% before and after Beagle imputation, respectively, for variants called from both assemblies. We observed only slight differences in the concordance metrics between variants called from either ARS-UCD1.2 or UOA_Angus_1, indicating that the genotypes of the 112 BSW cattle were accurately called from both assemblies, and that Beagle phasing and imputation further increased the genotyping accuracy.

**Table 2.** Comparisons between array-called and sequence variant genotypes. Non-reference sensitivity (NRS), non-reference discrepancy (NRD) and the concordance (CONC) between array-called and sequence-called genotypes for 112 BSW cattle that had BovineHD and sequence-called genotypes at 530,372 autosomal SNPs.

|  | GATK hard filtering |  |  | GATK hard filtering + Beagle imputation |  |  |
| --- | --- | --- | --- | --- | --- | --- |
|  | NRS | NRD | CONC | NRS | NRD | CONC |
| ARS-UCD1.2 | 99.14 | 2.75 | 98.13 | 99.77 | 0.60 | 99.59 |
| UOA_Angus.1 | 99.37 | 2.45 | 98.09 | 99.88 | 0.47 | 99.64 |

Because Beagle phasing and imputation improved the genotype calls from GATK, the subsequent analyses are based on the imputed sequence variant genotypes. After imputation, 81,674 (0.36%, 72,121 SNPs, 9,553 INDELs) and 104,217 (0.46%, 75,342 SNPs, 28,875 INDELs) variants were fixed for the alternate allele in ARS-UCD1.2 and UOA_Angus_1, respectively (Supplementary File 3: Table S3). Both the number and the percentage of variants fixed for the alternate allele was higher (0.10 percent points the latter, P = 0.027) for the UOA_Angus_1 than the ARS-UCD1.2 assembly. While the proportion and number of SNPs fixed for the alternate allele did not differ significantly (P = 0.65) between the assemblies, 0.61 percent points more INDELs (P = 1.45 × 10^−9^) were fixed for the alternate allele in UOA_Angus_1 than ARS-UCD1.2. 22,488,261 and 22,289,905 variants were polymorphic (*i.e.*, minor allele count *≥* 1) among the 161 BSW animals in ARS-UCD1.2 and UOA_Angus_1, respectively (Table 3). The number of variants detected per sample ranged from 6.91 to 8.58 million (7.28 ± 0.15) in ARS-UCD1.2 and from 6.93 to 8.44 million (7.26 ± 0.15) in UOA_Angus_1. More SNPs and INDELs were discovered for the ARS-UCD1.2 than UOA_Angus_1 assembly.

**Table 3.** Variants segregating among 161 BSW samples. Number of high-quality non-fixed variants discovered after aligning the samples to ARS-UCD1.2 and UOA_Angus_1 assemblies. Numbers in parentheses reflect the variant density (number of variants per Kb) along the autosomes.

|  | ARS-UCD1.2 | UOA_Angus.1 |
| --- | --- | --- |
| Non-fixed variants (per Kb) | 22,488,261 (9.03) | 22,289,905 (9.03) |
| Non-fixed SNPs (per Kb) | 19,557,039 (7.86) | 19,446,648 (7.88) |
| Non-fixed INDELs (per Kb) | 2,931,222 (1.18) | 2,843,257 (1.15) |

To take the length of the autosomes into consideration, we calculated the number of variants per Kb. While the overall variant and INDEL density was slightly higher for the ARS-UCD1.2 assembly, the SNP density was slightly higher for the UOA_Angus_1 assembly (Table 3).

The number and density of high-quality variants segregating on the 29 autosomes was 2.04 (P = 0.51) and 0.45 (P = 0.39) percent points higher, respectively, for the ARS-UCD1.2 than the UOA_Angus_1 assembly (Fig. 1, Supplementary File 4: Fig. S1). The difference in the number of variant sites detected from both assemblies was lower for SNPs (1.71 percent points) than INDELs (4.28 percent points). Chromosomes 9 and 12 were the only autosomes for which more variants were detected using the UOA_Angus_1 than ARS-UCD1.2 assembly. Differences in the number of variants detected were evident for chromosomes 12 and 28. While chromosome 12 has 29% more variants when aligned to UOA_Angus_1, chromosome 28 has 31% more variants when aligned to ARS-UCD1.2.

**Figure 1.**
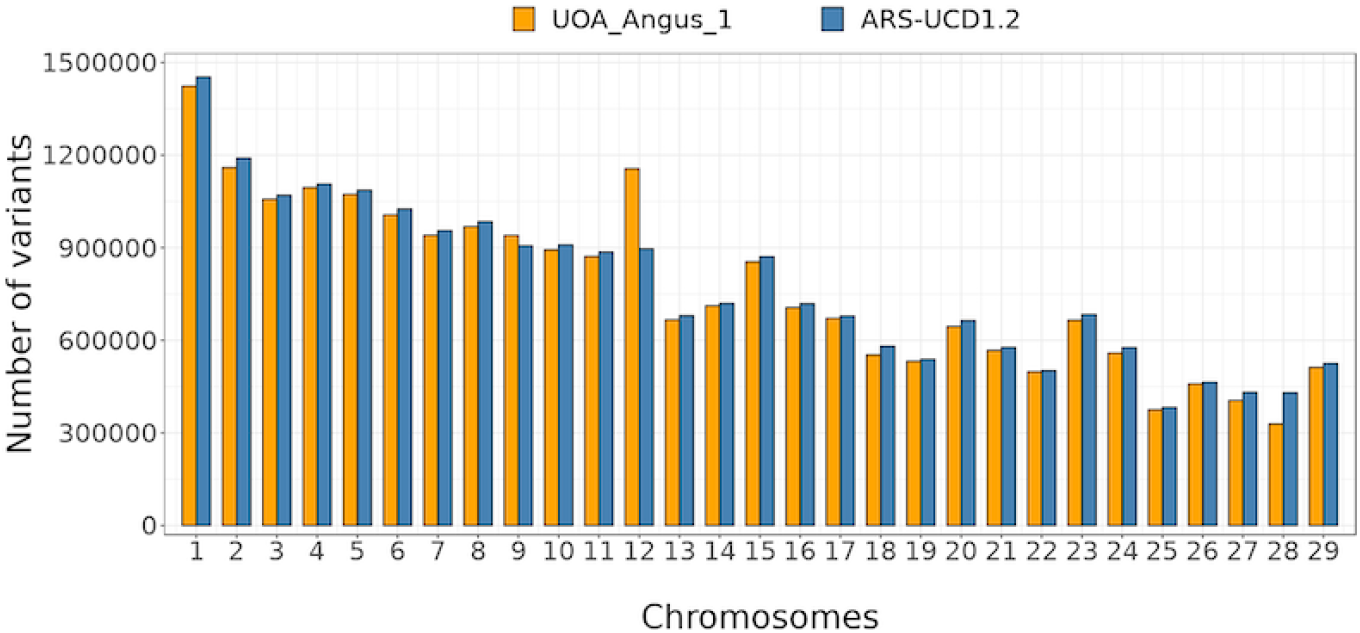
Total number of variants of autosomes for both assemblies. Number of variants detected on autosomes when the 161 BSW samples are aligned to the ARS-UCD1.2 (blue) and UOA_Angus_1 (orange) assembly.

The variant density of 26 out of the 29 autosomes (except for chromosomes 9, 12 and 26) was higher for the ARS-UCD1.2 assembly than the UOA_Angus_1 assembly. However, the density of INDELs was only higher for chromosome 12. Chromosome 23 had a higher variant density than all other chromosomes for both assemblies, with an average number of 13 variants detected per Kb. The high variant density at chromosome 23 primarily resulted from an excess of polymorphic sites within a ~5 Mb segment (between 25 and 30 Mb in the ARS-UCD1.2 and between 22 and 27 Mb in UOA_Angus_1) encompassing the bovine major histocompatibility complex (BoLA) (Supplementary File 5: Fig. S2). Other autosomes with density above 10 variants per Kb for both assemblies were chromosomes 12, 15 and 29. We observed the least variant density (~8 variants per Kb) at chromosome 13. Chromosome 12 carries a segment with an excess of variants at ~70 Mb in both assemblies. Visual inspection revealed that the segment with an excess of polymorphic sites was substantially larger in UOA_Angus_1 (7.6 Mb) than ARS-UCD1.2 (3.5 Mb) (Fig. 2). The variant-rich region at chromosome 12 coincides with a large segmental duplication that compromises reference-guided variant genotyping from short-read sequencing data and that has been described earlier [31, 32, 33]. Because of the greater number of variants and variant density in UOA_Angus_1, this extended region had a large impact on the cumulative genome-wide metrics presented in Table 3. When the same metrics were calculated without chromosome 12, the average density of both SNPs and INDELs was higher for ARS-UCD1.2 than UOA_Angus_1 (Supplementary File 6: Table S4). Segments with an excess of polymorphic sites were also detected on the ARS-UCD1.2 chromosomes 4 (113-114 Mb), 5 (98-105 Mb), 10 (22-26 Mb), 18 (60-63 Mb), and 21 (20-21 Mb). The corresponding regions in the UOA_Angus_1 assembly showed the same excess of polymorphic sites. However, these regions were shorter, and their variant density was lower compared to the extended segment at chromosome 12. The strikingly higher number (+31%) of variants discovered at chromosome 28 for ARS-UCD1.2 than UOA_Angus_1 was due to an increased length of chromosome 28 in the ARS-UCD1.2 assembly (Fig. 2).

**Figure 2.**
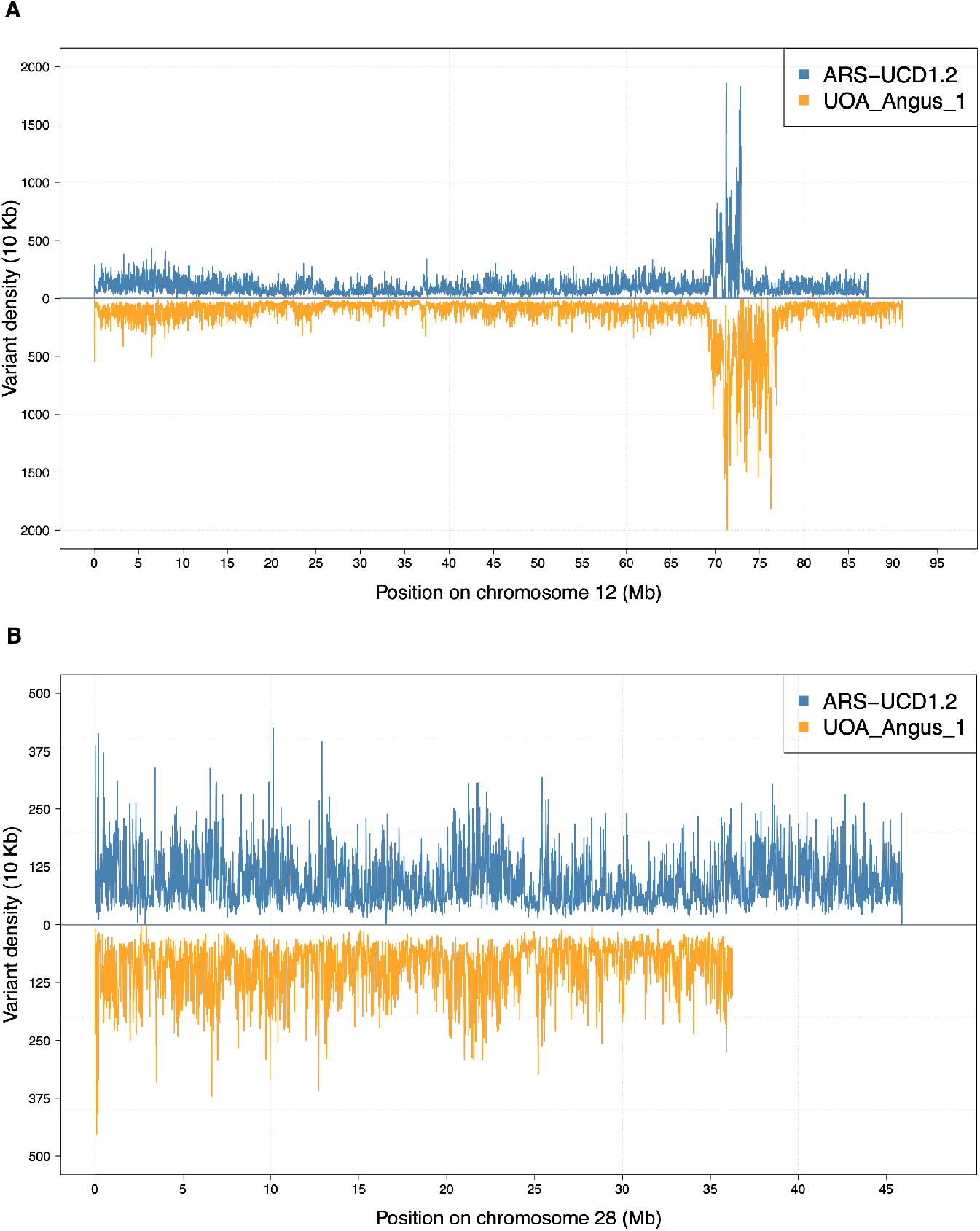
Density of variants across chromosomes 12 and 28. The number of variants within non-overlapping windows of 10 Kb for chromosome 12 (A) and 28 (B). The x-axis indicates the physical position along the chromosome (in Mb). The number of variants within each 10 Kb window is shown on the y-axis. Assembly ARS-UCD1.2 is displayed above the horizontal line (blue) and assembly UOA_Angus_1 is displayed below the horizontal line (orange).

Of 22,488,261 and 22,289,905 high-quality non-fixed variants, 848,100 (3.78%) and 857,206 (3.83%) had more than two alleles in the ARS-UCD1.2 and UOA_Angus_1 alignments, respectively (Supplementary File 7: Table S5). Most (69.75% for ARS-UCD1.2 and 69.09% for UOA_Angus_1) of the multi-allelic sites were INDELs. The difference in the percentage of multiallelic SNPs across assemblies was negligible. However, the difference in percentage of multiallelic INDELs was 0.69 percent points higher (P = 2.55 × 10^−9^) for UOA_Angus_1 than ARS-UCD1.2 autosomes.

In order to detect potential flaws in sequence variant genotyping, we investigated if the genotypes at the high-quality non-fixed variants agreed with Hardy-Weinberg proportions. We observed 218,734 (0.97%) and 243,408 (1.09%) variants for ARS-UCD1.2 and UOA_Angus_1, respectively, for which the observed genotypes deviated significantly (P *<* 10^−8^, Supplementary File 7: Table S5) from expectations. The proportion of high-quality non-fixed variants for which the genotypes do not agree with Hardy-Weinberg proportions is 0.12 percent points higher for the UOA_Angus_1 than ARS-UCD1.2 assembly. At chromosome 12, 3.29 percent points more variants deviated from Hardy-Weinberg proportions for the UOA_Angus_1 than the ARS-UCD1.2 assembly (Supplementary File 8: Fig. S3); more than twice the difference observed for any other autosome. When variants located on chromosome 12 were excluded from this comparison, we observed 199,304 (0.92%) and 180,264 (0.85%) variants for the ARS-UCD1.2 and UOA_Angus_1 assembly, respectively, for which the observed genotypes deviated significantly (P *<* 10^−8^) from expectations.

### Functional annotation of polymorphic sites

Using the VEP software, we predicted functional consequences based on the Ensembl genome annotation for 19,557,039 and 19,446,648 SNPs, and 2,931,222 and 2,843,257 INDELs, respectively, that were discovered from the ARS-UCD1.2 and UOA_Angus_1 alignments. Most SNPs were in either intergenic (66.30% and 56.56%) or intronic regions (32.55% and 42.09%) for ARS-UCD1.2 and UOA_Angus_1, respectively (Table 4, Supplementary File 9: Table S6). Only 224,549 and 262,775 (1.15% and 1.35%) of the SNPs were in exons for ARS-UCD1.2 and UOA_Angus_1, respectively. The majority of INDELs was in either intergenic (65.76% and 55.95%) or intronic regions (33.84% and 43.47%) for ARS-UCD1.2 and UOA_Angus_1, respectively (Table 4, Supplementary File 9: Table S6). Only 11,561 and 16,391 (0.40% and 0.58%) INDELs were in exonic sequences. While the number and proportion of variants in coding regions was similar for both assemblies, we observed marked differences in the number of variants annotated to intergenic and intronic regions. The percentage of SNPs and INDELs annotated to intergenic regions is 9.74 and 9.81 percent points higher, respectively, for the ARS-UCD1.2 than UOA_Angus_1 assembly. In contrast, the percentage of SNPs and INDELs annotated to intronic regions is 9.54 and 9.63 percent points higher, respectively, for the UOA_Angus_1 than the ARS-UCD1.2 assembly. According to the Ensembl annotation of the autosomal sequences, intergenic, intronic and exonic regions span respectively 61.53, 34.77 and 3.80% in ARS-UCD1.2 and 52.32, 42.32 and 5.36% in UOA_Angus_1.

**Table 4.** Number of SNPs and INDELs annotated using the VEP software per region and assembly. Annotated SNPs and INDELs are classified by region were detected. The total number of annotated variants per assembly and region are displayed here. The table lists only the most severe annotation. The percentage of variants placed in each region per variant type and assembly is shown between parentheses.

|  | ARS-UCD1.2 |  | UOA_Angus_1 |  |
| --- | --- | --- | --- | --- |
|  | SNPs | INDELs | SNPs | INDELs |
| Exonic regions (%) | 224,549 (1.15) | 11,561 (0.40) | 262,775 (1.35) | 16,391 (0.58) |
| Intronic regions (%) | 6,365,765 (32.55) | 992,015 (33.84) | 8,185,503 (42.09) | 1,236,006 (43.47) |
| Intergenic regions (%) | 12,966,725 (66.30) | 1,927,646 (65.76) | 10,998,370 (56.56) | 1,590,860 (55.95) |

Either moderate or high impacts on protein function were predicted for 89,812 and 103,576 SNPs, and 10,259 and 11,847 INDELs (0.46 and 0.53% of the total annotated SNPs and 0.35 and 0.41% of the total annotated INDELs), respectively, that were discovered from ARS-UCD1.2 and UOA_Angus_1 alignments (Tables 5 and 6).

**Table 5.** SNPs in high or moderate effect categories. Number of SNPs in high and moderate (marked with an asterisk) effect categories per assembly.

|  | ARS-UCD1.2 | UOA_Angus_1 |
| --- | --- | --- |
| Missense variant* | 86,634 | 99,773 |
| Stop gained | 1,466 | 1,911 |
| Splice donor variant | 506 | 525 |
| Splice acceptor variant | 345 | 395 |
| Start lost | 271 | 319 |
| Stop lost | 155 | 218 |

**Table 6.** INDELs in high or moderate effect categories. Number of INDELs in high and moderate (marked with an asterisk) effect categories per assembly.

|  | ARS-UCD1.2 | UOA_Angus_1 |
| --- | --- | --- |
| Frameshift variant | 6,289 | 7,435 |
| Inframe deletion* | 1,761 | 1,972 |
| Inframe insertion* | 850 | 985 |
| Splice donor variant | 291 | 298 |
| Splice acceptor variant | 292 | 292 |
| Stop gained | 218 | 288 |
| Protein altering variant* | 87 | 107 |
| Start lost | 20 | 14 |
| Stop lost | 11 | 15 |
| Transcript ablation | 5 | 6 |

The number of variants with putatively high or moderate effects was higher for the UOA_Angus_1 than ARS-UCD1.2 assembly for 14 of 16 functional classes of annotations. Differences across all autosomes were observed for SNPs that potentially affect splice acceptor variants (345 for ARS-UCD1.2 and 395 for UOA_Angus_1, P = 0.032) and SNPs that potentially cause the loss of a stop codon (155 for ARS-UCD1.2 and 218 for UOA_Angus_1, P = 0.037). Differences across all autosomes also resulted for INDELs that potentially cause inframe deletions (1,761 for ARS-UCD1.2 and 1,972 for UOA_Angus_1, P = 0.0035), INDELs that potentially cause inframe insertions (850 for ARS-UCD1.2 and 985 for UOA_Angus_1, P = 0.0013) and INDELs that potentially cause the gain of a stop codon (218 for ARS-UCD1.2 and 288 for UOA_Angus_1, P = 0.016).

### Signatures of selection

Next, we investigated how the choice of the reference genome impacts the detection of putative signatures of selection in the 161 BSW cattle. We used the composite likelihood ratio (CLR) test to identify beneficial adaptive alleles that are either close to fixation or recently reached fixation [34]. As information on ancestral and derived alleles was not available, we considered 19,370,683 (ARS-UCD1.2) and 19,255,155 (UOA_Angus_1) sequence variants that were either polymorphic or fixed for the alternate allele in the 161 BSW cattle. The CLR test revealed 40 and 33 genomic regions (merged top 0.1% windows) encompassing ~2.5 and ~2.48 Mb, and 29 and 27 genes, respectively, from the ARS-UCD1.2 and the UOA_Angus_1 alignments (Fig. 3, Supplementary File 10: Table S7, Supplementary File 11: Table S8).

**Figure 3.**
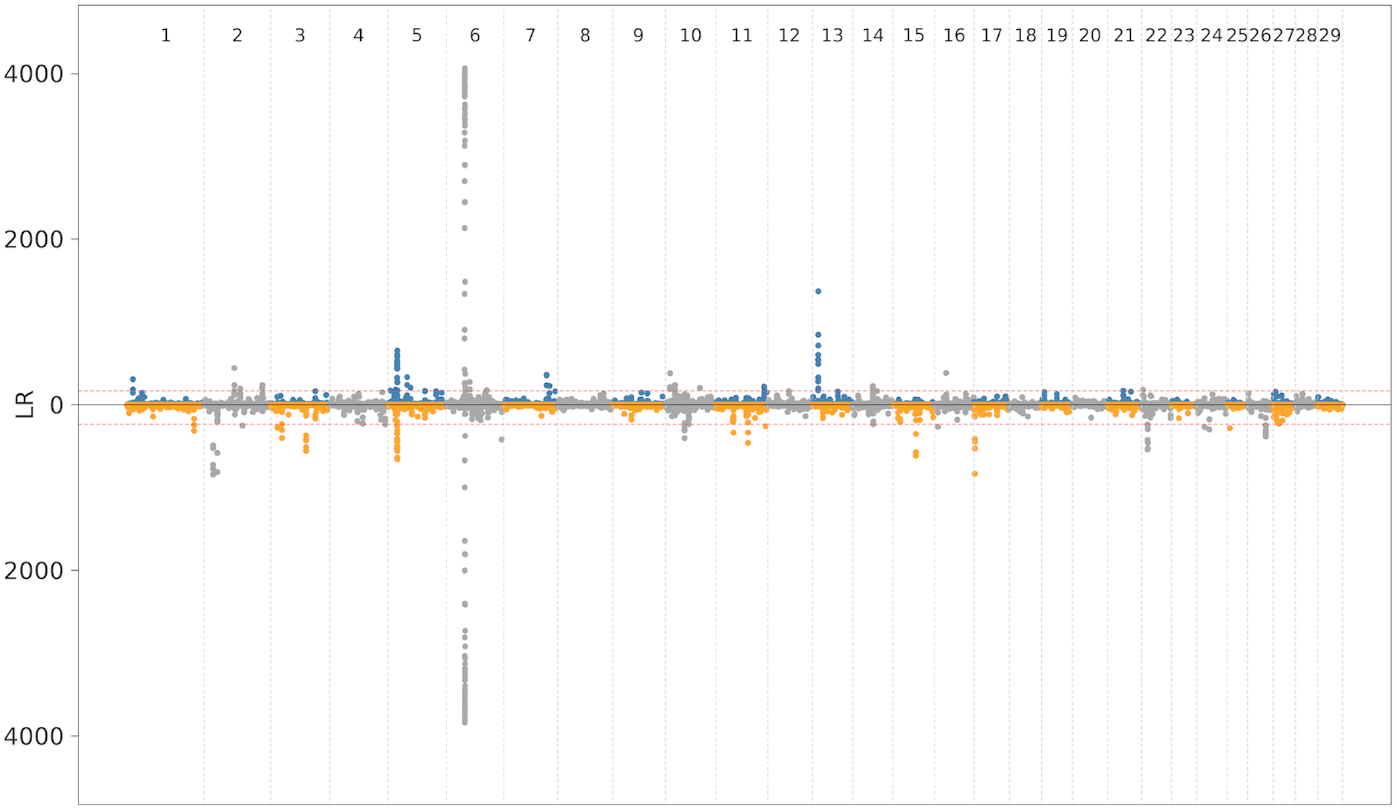
Genome wide distribution of selection signals from CLR. Selection signal distribution for both ARS-UCD1.2 (top panel) and UOA_Angus_1 assemblies (bottom panel). Red dotted line shows top 0.1% signal.

A putative signature of selection on chromosome 6 encompassing the *NCAPG* gene had high CLR values in both assemblies (CLR_ARS-UCD1.2_ = 4064; CLR_UOA_Angus_1_ = 3838). Another signature of selection was detected for both assemblies upstream the *KITLG* gene on chromosome 5 (ARS-UCD1.2: 18.48 - 18.86 Mb, CLR_ARS-UCD1.2_ = 655; UOA_Angus_1: 18.48 - 18.84, CLR_UOA_Angus_1_ = 657). However, most of the signatures of selection were detected for only one assembly. A putative selective sweep on chromosome 13 was identified using the ARS-UCD1.2 but not the UOA_Angus_1 assembly. The putative selective sweep was between 11.5 and 12 Mb encompassing three protein coding (*CCDC3*, *CAMK1D* and ENSBTAG00000050894) and one non-coding gene (ENSBTAG00000045070). The top window (CLR=1373) was between 11,962,310 and 12,022,317 bp. In order to investigate why the CLR test revealed strong evidence for the presence of a signature of selection in ARS-UCD1.2 but not in UOA_Angus_1, we investigated the corresponding region in both assemblies using dot plots, variant density, alternate allele frequency and alignment coverage. The dot plot revealed that the orientation of bovine chromosome 13 is flipped in the UOA_Angus_1 assembly. The putative signature of selection is next to but clearly distinct from a region with a very high SNP density and sequence coverage in both assemblies (Supplementary File 12: Fig. S4). We detected 350 SNP within the top window (5.87 SNP / Kb) of which 145 were fixed for the alternate allele. Within the corresponding region on UOA_Angus_1, we detected 209 SNP (3.48 SNP / Kb) of which 13 were fixed for the alternate allele. This pattern indicates that the 161 sequenced BSW cattle carry a segment in the homozygous state that is more similar to the UOA_Angus_1 than ARS-UCD1.2 reference genome. We observed the reciprocal pattern for a putative selective sweep on chromosome 22 that was detected using UOA_Angus_1 but not ARS-UCD1.2 (Supplementary File 13: Fig. S5).

### Genome-wide association testing

Next, we imputed genotypes for autosomal variants that were detected using the two assemblies for 30,499 cattle that had (partially imputed) Illumina BovineHD array-derived genotypes. The average imputation accuracy (Beagle R^2^) was 0.87 ± 0.27 (median: 0.99) in the ARS-UCD1.2 and 0.87 ± 0.26 (median: 0.99) in the UOA_Angus_1 assembly. To prevent bias resulting from imputation errors, we removed variants that had low frequency (minor allele count *<* 3), low accuracy of imputation (Beagle R^2^ *<* 0.5) or for which the observed genotypes deviated significantly (P *<* 10^−6^) from Hardy-Weinberg proportions from the imputed data. Following quality control, 12,761,165 and 12,602,069 imputed variants were respectively retained (with imputation accuracy of 0.95 ± 0.11 and 0.95 ± 0.10) for genetic investigations in the ARS-UCD1.2 and UOA_Angus_1 dataset representing 56.75 and 56.54% of the 22,488,261 and 22,289,905 high-quality segregating variants. We then carried out genome-wide association studies (GWAS) between imputed sequence variant genotypes and six traits, including stature and five dairy traits (milk yield, fat yield, protein yield, protein and fat percentage), for which between 11,294 and 12,396 cattle had phenotypes in the form of de-regressed proofs. The resulting Manhattan plots appeared very similar for both datasets (Fig. 4, Supplementary File 14: Fig. S6). Across the six traits analysed, the number of significantly associated variants was similar when the association analyses were performed using imputed sequence variants identified in the two builds. The difference in the number of significantly associated variants (P *<* 10^−8^) between the two builds is mainly due to variants that had P-values that were slightly above the threshold of 10^−8^ in one but not the other build.

**Figure 4.**
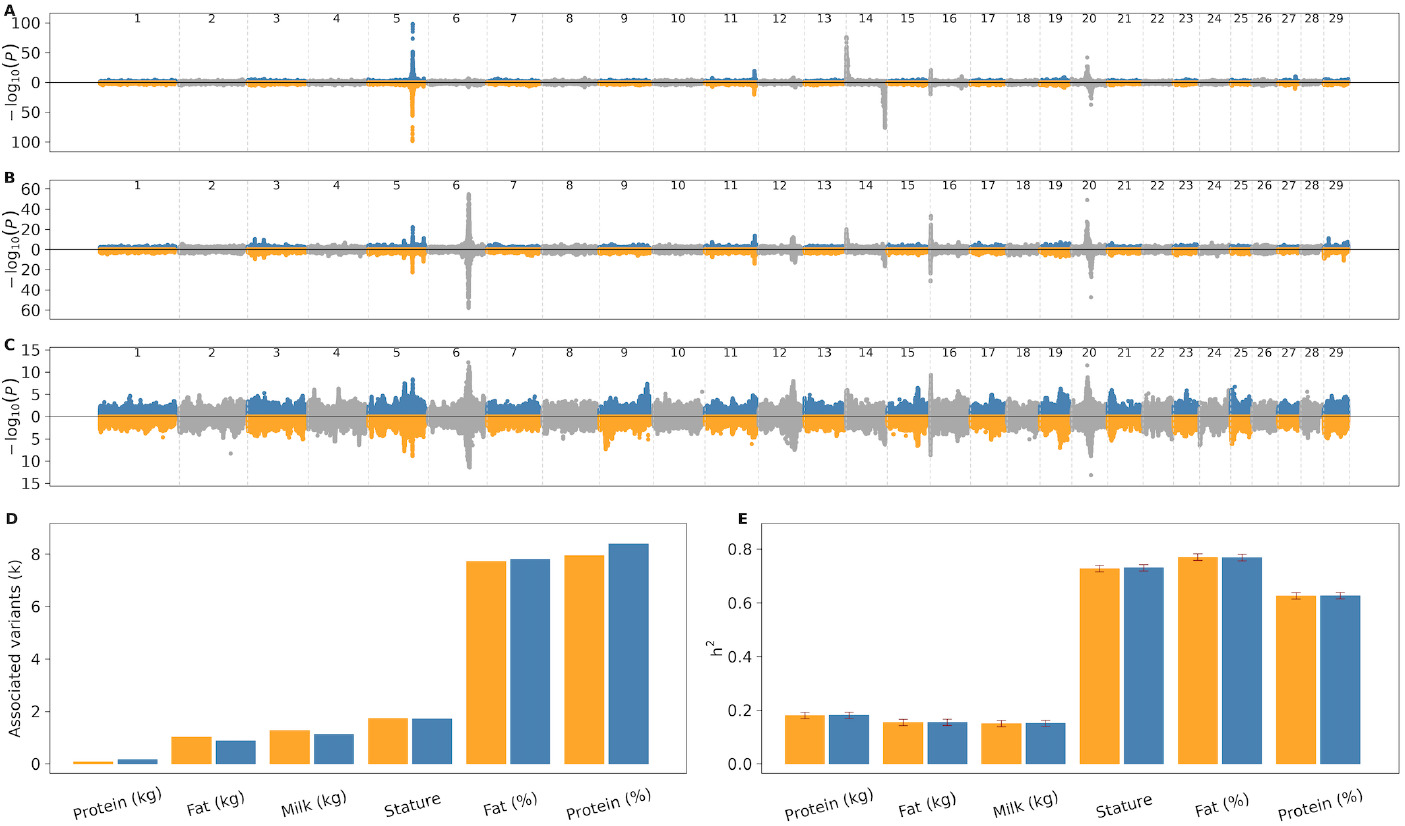
Manhattan plots for fat percentage, protein percentage, and milk yield. Number of significantly (P *<* 10-8) associated variants in GWAS for seven traits. Estimated genomic heritability for stature and six dairy traits. Manhattan plots showing association of sequence variants - imputed using ARS-UCD1.2 (blue and grey) and UOA_Angus_1 (orange and grey) - with fat percentage (A), protein percentage (B) and milk yield (C). The orientation of some autosomes (*e.g.*, chromosome 14) is flipped between ARS-UCD1.2 and UOA_Angus_1. The number (in thousands) of variants - imputed using ARS-UCD1.2 (blue) and UOA_Angus_1 (orange) - significantly (P *<* 10^−8^) associated with the seven traits considered for GWAS (D). Genomic heritability estimated using all autosomal variants imputed using ARS-UCD1.2 (blue) and UOA_Angus_1 assemblies (orange) (E). Standard errors of the estimates are indicated in read lines.

To investigate if causal variants can be readily identified from both assemblies, we inspected the QTL for dairy traits at chromosomes 14 and 20, respectively, for which p.Ala232Lys in *DGAT1* encoding Diacylglycerol O-Acyltransferase 1 and p.Phe279Tyr in *GHR* encoding Growth Hormone Receptor have been proposed as causal variants [35, 36]. The accuracy of imputation for the Phe279Tyr variant in the *GHR* gene was 0.92 and 0.88 for the ARS-UCD1.2 and UOA_Angus_1 assembly, respectively. In the association studies for milk yield, fat percentage and protein percentage, for which chromosome 20 QTL was detected, the p.Phe279Tyr variant was the most significantly associated variant in both assemblies. The SNP is located at 31,888,449 and 39,903,176 bp on the ARS-UCD1.2 and UOA_Angus_1 build. The frequency of the milk yield-increasing and fat and protein content-decreasing tyrosine-encoding T allele was 12.90 and 13.02% in ARS-UCD1.2 and UOA_Angus_1, respectively, and the P-values for milk yield, fat percentage and protein percentage were 3.18 × 10^−12^, 1.11 × 10^−42^, 6.98 × 10^−50^ and 7.40 × 10^−14^, 6.89 × 10^−38^, 5.57 × 10^−48^.

Two adjacent SNPs (ARS-UCD1.2: g.611019G*>*A & g.611020C*>*A; UOA_Angus_1: g.81672806C*>*T & g.81672805G*>*T), in the coding sequence of *DGAT1* cause the p.Ala232Lys substitution that has a large effect on milk yield and composition. In 161 sequenced BSW cattle of our study, the alternate allele was detected in the heterozygous state in two and one animals using the ARS-UCD1.2 and UOA_Angus_1 datasets. When imputed into array-derived genotypes of the mapping cohort, the lysine variant had a frequency of 0.0082 (Beagle R^2^: 0.98) and 0.0002 (Beagle R^2^: 0.82) in the ARS-UCD1.2 and UOA_Angus_1 imputed genotypes. An association study between imputed sequence variant genotypes and fat percentage revealed strong association (P = 1.46 × 10^−76^) at the proximal region of chromosome 14 encompassing *DGAT1* in the ARS-UCD1.2 data (Fig. 4A). The top association signal resulted from a variant at position 420,486. The P-value of the p.Ala232Lys variant was only slightly higher (P = 2.18 × 10^−76^). Using the UOA_Angus_1 imputed data, we detected strong association at the corresponding region (Fig. 4A). The most significantly associated variant (P = 1.80 × 10^−76^) was at 81,673,955 bp. However, the p.Ala232Lys variant was not associated with fat percentage (P = 0.33). Also, the *DGAT1* gene was missing in the Ensembl annotation of the UOA_Angus_1 assembly.

Next, we estimated the genomic heritability (h^2^) for stature and six dairy traits using a genomic restricted maximum likelihood estimation (GREML) approach. Therefore, we built a genomic relationship matrix separately for each assembly using the genotypes of all imputed autosomal variants that had minor allele count *>* 3 and imputation accuracy (Beagle R^2^) *>* 0.5. The estimates for the genomic h^2^ did not differ for all seven traits (Fig. 4E). We then partitioned (genomic) h^2^ by the 29 autosomes using the two imputed datasets. As seen for the total h^2^, we found no difference in variance explained by individual autosomes between the two assemblies.

## Discussion

We investigated whether the choice of the reference genome impacts genomic analyses in BSW cattle that have been sequenced with short paired-end reads. To the best of our knowledge, such an evaluation had not been performed so far in cattle. A Hereford-based genome assembly [20] is accepted by the bovine genomics community as reference genome for reference-guided alignment and variant detection in both taurine and indicine cattle [8, 9]. Recently, the application of sophisticated methods to assemble long sequencing reads provided reference-quality assemblies for cattle breeds other than Hereford [17, 21]. None of these novel reference-quality assemblies has been considered as a reference genome for sequence variant analysis so far. The genetic distance between the reference genome and the target sample and the properties (GC content, genome size, proportion of repeats) of the reference genome impact reference-guided mapping and variant genotyping [17, 24, 25, 37, 38]. To investigate reference-guided sequence analyses from different assemblies, we aligned short sequencing reads of 161 BSW cattle to the Hereford-based ARS-UCD1.2 and Angus-based UOA_Angus_1 assemblies. Widely used metrics (contig N50, scaffold N50, BUSCO completeness) suggest that both assemblies are of reference quality [17, 20]. The sequence read mapping and variant genotyping accuracy did not differ notably between the ARS-UCD1.2 and UOA_Angus_1 assemblies, indicating that both assemblies are suitable for reference-guided genome analyses in BSW cattle. The BSW, Angus and Hereford breeds are closely related as these breeds diverged relatively recently [39]. Greater genetic distance between the target breed and the reference genome might compromise mapping rate and alignment quality [24, 25, 40]. However, it is worth mentioning that the orientation of some chromosomes is flipped in UOA_Angus_1 (*i.e.*, the beginning of the chromosome corresponds to the end in the corresponding ARS-UCD1.2 entry). This does not affect sequence read mapping and variant genotyping but needs to be considered when comparing selection signatures and association signals across assemblies.

The number and density of INDELs that segregate in 161 BSW cattle was slightly lower when variants were called from the UOA_Angus_1 than ARS-UCD1.2 alignment. However, the proportion of multiallelic INDELs and INDELs fixed for the alternate allele was higher in the UOA_Angus_1 than ARS-UCD1.2 alignment. In fact, the absolute number of INDELs fixed for the alternate allele was three times higher when the sequence data were aligned against the UOA_Angus_1 assembly. An excess of artefactual INDELs in long-read sequencing-based assemblies was noted by Watson and Warr [41]. Both the ARS-UCD1.2 and UOA_Angus_1 assembly were constructed from PacBio continuous long reads. While ARS-UCD1.2 was polished with short reads and manually curated, this step was not as extensively carried out for the UOA_Angus_1 assembly [17, 20]. Our results may indicate that UOA_Angus_1 contains somewhat more artefactual INDELs than ARS-UCD1.2. However, the absolute number of artefactual INDELs is low for both assemblies and their genotypes are likely to be discarded from most downstream analyses as most of them will be fixed for the alternate allele. Importantly, the concordance between sequence- and array-called genotypes was very high and the number of variants deviating from Hardy-Weinberg proportions was very low at segregating sites for both assemblies, indicating that reliable genotypes can be obtained from both ARS-UCD1.2 and UOA_Angus_1.

The length of chromosomes 12 and 28 differs considerably between the assemblies. A large segmental duplication affects chromosome 12 in both assemblies. This duplication compromises the mapping of sequencing reads, thereby causing misalignments and flaws in the resulting genotypes [31, 32, 33]. An excess of variants, including many for which the genotypes deviate from Hardy-Weinberg proportions, was detected for both assemblies within the segmental duplication. Because the segmental duplication is two times longer in UOA_Angus_1 than ARS-UCD1.2, the genome-wide number of variants, variant density, proportion of missing genotypes and number of variants deviating from Hardy-Weinberg proportions was higher using UOA_Angus_1. At chromosome 28, the variant density was similar for both assemblies, but the absolute number of variants detected was lower for UOA_Angus_1 because the chromosome was shorter. The UOA_Angus_1 assembly lacks approximately 9.5 million bases that likely correspond to the ARS-UCD1.2 chromosome 28 sequence from 36,496,661 bp onwards. According to the Ensembl (build 101) annotation of ARS-UCD1.2, this segment encompasses 67 genes that are consequently missing in the autosomal annotation of UOA_Angus_1.

Differences in the functional annotations predicted for variants obtained from ARS-UCD1.2 and UOA_Angus_1 were evident from the output of the VEP tool. The number of variants annotated to inter- and intragenic regions differed between the assemblies because the length of these features differed in the annotation files. An example for a striking difference in the coding sequence between both annotations is *DGAT1*, a gene that harbours a missense variant (p.Ala232Lys) with a large impact on dairy traits [36]. Our GWAS identified a QTL for dairy traits at chromosome 14 in both assemblies. The QTL encompassed *DGAT1* using the ARS-UCD1.2 annotation. However, *DGAT1* was not annotated at the corresponding sequence of the UOA_Angus_1 assembly. Given the manual curation efforts of the ARS-UCD1.2 annotation in contrast to the mere computational-based inference of annotations for UOA_Angus_1 from the Ensembl database, we suspect that the latter produces more erroneous annotations [42]. In fact, the ARS-UCD1.2 assembly is currently the widely accepted and universally applied bovine reference genome [43, 44]. It is very unlikely that this will change soon because besides the completeness and continuity of the reference assembly, its functional annotation is crucial for downstream analyses. While tools exist to lift physical coordinates from one genomic context to another based on flanking sequences, this approach is cumbersome. Consequently, errors and gaps in the functional annotations of bovine reference-quality assemblies other than ARS-UCD1.2 are a major obstacle to switch references. TThe application of an augmented reference genome that contains ARS-UCD1.2 and its functional annotations as backbone as well as variants detected in other assemblies might solve such problems [12, 45].

We applied the composite likelihood ratio test to detect alleles that are either close to fixation or already reached fixation using genotypes obtained from both references. Supplying information about ancestral and derived alleles to the composite likelihood ratio test is required to determine which allele has been under selection and increases the statistical power to detect signatures of selection [34, 46]. Although we were unable to differentiate between ancestral and derived alleles, we identified strong signatures of selection from both assemblies at regions that were previously detected in different cattle breeds including BSW [47, 48, 49]. This finding suggests that plausible signatures of selection can be identified using folded site frequency spectrum. However, most signatures of selection did not overlap between both assemblies. For instance, a strong selective sweep on chromosome 13 was only detected using the ARS-UCD1.2 assembly, while a putative sweep on chromosome 22 was only detected using the UOA_Angus_1 assembly. These differences were unexpected because the two assemblies were constructed from closely related breeds. In fact, Hereford and Angus are both taurine beef breeds that are genetically closely related [39]. When the BSW samples were aligned to the ARS-UCD1.2 assembly, the chromosome 13 region harbouring the signature of selection was depleted for variation, suggesting that the selected allele(s) already reached fixation. In fact, we observed many variants that were fixed for the alternate allele within the top windows at chromosome 13. These variants were absent when the sequencing data were aligned to UOA_Angus_1, because their alternate alleles in ARS-UCD1.2 correspond to reference alleles in UOA_Angus_1. Thus, our findings suggest that detecting selective sweeps that already reached fixation with the composite likelihood ratio test depends on the relationship between the study population and the reference genome if a folded site frequency spectrum is used. The CLR test would reveal the same regions from both assemblies if only segregating sites are considered for the analysis. However, restricting the analysis to segregating sites bears a risk of missing sweeps that already reached fixation.

To our knowledge, a quantitative assessment of differences arising from the use of different reference genomes had only been performed in humans at a single nucleotide variant (SNV) level [25, 38]. Recently, Low *et al.* [17] mapped 38 cattle samples from 7 breeds against the Brahman and Angus assemblies to detect larger structural variants that may be involved in the adaptability of indicine cattle to harsh environments. We considered 161 BSW cattle for a thorough characterization of reference-guided analyses from two assemblies. As such an evaluation may be regularly performed in the future for many species, we developed a workflow that can be adapted and reused for varies breeds, populations and species [50]. In fact, our evaluation is the first to compare sequence variant discovery from primary and haplotype-resolved assemblies. Therefore, our findings also show that haplotype-resolved reference-quality assemblies may readily serve as reference genomes for linear read mapping and variant genotyping.

## Conclusions

Our results suggest that both the ARS-UCD1.2 and UOA_Angus_1 assembly are suitable for reference-guided genome analyses in BSW cattle. The choice of the reference may have a large impact on detecting signatures of selection that already reached fixation. Furthermore, curation of the reference genomes is required to improve the characterisation of functional elements. The workflow herein developed is a starting point for a comprehensive comparison of the impact of reference genomes on genomic analyses in various breeds, populations and species.

## Methods

### Data availability and code reproducibility

Short paired-end whole-genome sequencing reads of 161 BSW cattle were considered for our analyses. Accession numbers for all animals are available in the Supplementary File 1: Table S1.

In order to investigate the effect of different assemblies on downstream analyses, we considered the current bovine Hereford-based reference genome (ARS-UCD1-2) [20] and an Angus-based reference-quality assembly (UOA_Angus_1) [17] that was generated from a F1 Angus x Brahman cross. The assemblies were downloaded from the public repositories of the NCBI (GCA 002263795.2, GCA 003369685.2). The UOA_Angus_1 assembly does not contain the X chromosomal sequence because it represents the paternal haplotype of a male animal. The ARS-UCD1.2 assembly was created from a female cow, thus does not contain a Y chromosomal sequence. For the sake of completeness, we expanded the ARS-UCD1.2 assembly with the Y chromosomal sequence from Btau 5.0 and the UOA_Angus_1 assembly with the X chromosomal sequence from ARS-UCD1.2.

Alignment, coverage, variant calling, imputation, annotation and analysis workflows were implemented as described below using Snakemake [51] (version 5.10.0). Python 3.7.4 has been used for running custom scripts as well as for submission and generation of Snakemake workflows.

Unless stated otherwise, the R (version 3.3.3) software environment and ggplot2 package (version 3.0.0) were used to create figures and perform statistical analyses. Paired t-test and Kruskal-Wallis rank sum test were applied to assess differences between assemblies for normal and not normal distributed values, respectively.

### Alignment quality and depth of coverage

Quality assessment and control (removal of adapter sequences and reads and bases with low quality) of the raw sequencing data was carried out using the fastp software [52] (version 0.19.4) with default parameter settings. Reads were discarded when the phred-scaled quality was below 15 for more than 15% of the bases.

When necessary, the resulting FASTQ files were split into up to 13 read-group-specific FASTQ files to facilitate the read group aware processing of the data using gdc-fastq-splitter [53] (version 0.0.1). The filtered reads were subsequently aligned to both the ARS-UCD1.2 and UOA_Angus_1 assemblies (see above) using the MEM-algorithm of the Burrows-Wheeler Alignment (BWA) software [54, 55] (version 0.7.17) with option -M and –R to mark shorter split hits as secondary alignments and supply read group identifier and default values for all other parameters. Samblaster [56] (version 0.1.24) was used to mark duplicates in the SAM files, which were then converted into the binary format by using SAMtools [57] (version 1.6). Sambamba [58] (version 0.6.6) was used for coordinate-sorting (sort function) and to combine the read group-specific BAM files into sample-specific sorted BAM files. Duplicated reads and PCR duplicates of the merged and coordinate-sorted BAM files were marked using the MarkDuplicates module from Picard Tools [59] (version 2.18.17).

Uniquely mapped and properly paired reads that had mapping quality greater than 10 were obtained using SAMtools view -q 10 -F 1796. We considered a phredscaled mapping quality threshold of 10 to retain only reads (referred to as high-quality reads) that qualify for variant genotyping according to best practice guidelines of the GATK [28, 29].

The mosdepth software [60] (version 0.2.2) was used to extract the number of reads that covered a genomic position in order to obtain the average coverage per sample and chromosome. We considered only high-quality reads (by excluding reads with mapping quality *<* 10 and SAM flag 1796).

### Sequence variant genotyping and variant statistics

We used the BaseRecalibrator module of the Genome Analysis Toolkit (GATK - version 4.1.4.1) [61, 62] to adjust the base quality scores using 115,815,241 (ARS-UCD1.2) and 87,710,119 (UOA_Angus_1) unique positions from the Bovine dbSNP version 150, as known variants. To obtain the coordinates of known sites for the UOA_Angus_1 assembly, we used liftover coordinates obtained from the mapping of 120 bases flanking the known ARS-UCD1.2 positions to UOA_Angus_1 using the MEM-approach of BWA (see above) with option –k 120 to consider only full-length matches. To discover and genotype variants from the recalibrated BAM files, we used the GATK according to the best practice guidelines [28, 29]. The GATK HaplotypeCaller module was run to produce gVCF (genomic Variant Call Format) files. The gVCF files were then consolidated using GenomicsDBImport and passed to the GenotypeGVCFs module to genotype polymorphic SNP and INDELs. We applied the VariantFiltration module for site-level filtration with the following recommended thresholds to retain high-quality SNP and INDELs: QualByDepth (QD) *>* 2.0, Qual *>* 30, Strand Odds Ratio (SOR) *<* 3.0, FisherStrand (FS) *<* 60.0, RMSMappingQuality (MQ) *>* 40.0, MappingQualityRankSumTest (MQRankSum) *>* 12.5, ReadPosRankSumTest (ReadPosRankSum) *>* 8.0 for SNPs, and (QD) *>* 2.0, Qual *>* 30, Strand Odds Ratio (SOR) *<* 10.0, FisherStrand (FS) *<* 200.0, ReadPosRankSumTest (ReadPosRankSum) *>* −20.0 for INDELs. For both SNP and INDELs, the number of minimum alleles (AN) per variant was set up to the number of samples and variants not meeting all the criteria were discarded.

Beagle [30] (version 4.1) haplotype phasing and imputation was run to improve the raw genotype calls and impute missing genotypes. The genotype likelihood (gl) mode was applied in order to infer missing and adjust existing genotypes based on the phred-scaled likelihoods of all other non-missing genotypes.

Alternate allele frequency was calculated using the ‘–keep-allele-order –freq’ flags with PLINK 1.9 [63] and non-segregating variants were subsequently filtered out from the imputed VCF file with the option ‘–mac 1 –remove-filtered-all’ from VCFtools [64]. Biallelic variants have been retrieved by using the filter ‘–minalleles 2 –max-alleles 2’ with VCFtools. Index and stats for the relevant VCF files were generated through tabix [65], VCFtools and BCFtools [66], respectively. Per-sample stats were obtained by adding the ‘-v’ flag when generating the stats with VCFtools. Observed genotypes were tested for deviation from Hardy-Weinberg pro-portions using the ‘–hwe 10e-8’ and ‘–hardy –recode’ flags with PLINK 1.9 [63]. Transition and transversion ratio of SNPs were calculated via VCFtools.

### Functional annotation of polymorphic sites

Functional consequences of high-quality and non-fixed SNPs and INDELs were predicted according to the Ensembl (release 101) annotation of the bovine genome assembly ARS-UCD1.2 and UOA_Angus_1, respectively, using the Ensembl Variant Effect Predictor tool (VEP - version 91.3) [67] with default parameters and ‘–hgvs –symbol’ nomenclature. The classification of variants according to sequence ontology terms and the prediction of putative impacts on protein function followed Ensembl guidelines. Basic statistics of the annotation were calculated using AGAT [68] (version v0.5.1).

### Signatures of selection

Signatures of recent selection were identified using the composite likelihood ratio (CLR) approach implemented in Sweepfinder2 [69]. We considered 19,370,683 (ARS-UCD1.2) and 19,255,155 (UOA_Angus_1) biallelic SNP (segregating sites and SNP that were fixed for the non-reference allele) to calculate the CLR in 20 Kb windows with pre-computed empirical alternate allele frequency. The top 0.1% windows were considered as putative selective sweeps. Adjacent top 0.1% windows were merged into regions. The gene content of the regions was determined according to the annotations from Ensembl (release 101) using BEDTools [70].

### Dot plots

To identify sequence similarities and dissimilarities between the two assemblies, we inspected chromosome wise dot plots of pair-wise sequence alignments using LASTZ [71] (v1.04.03) with the options ‘–notransition –nogapped –step=20 –exact=50’ using repeat-masked assemblies which we downloaded from Ensembl (release 101).

### Imputation

Microarray-derived SNP genotypes were available for 30,499 BSW cattle typed on seven low-density (20k-150k) and one high-density chip (Illumina BovineHD; 777k). Coordinates of the SNP were originally determined according to the ARS-UCD1.2 build. To remap the SNP to the UOA_Angus_1 assembly, we used liftover coordinates obtained from the mapping of 120 bases flanking the BovineHD probes to the UOA_Angus_1 assembly using the MEM algorithm of BWA [54, 55] with option –k 120 to consider only full-length matches. Both the original and the remapped genotype data were imputed (separately) to the whole genome sequence level using a stepwise approach with reference panels aligned to the respective genome assemblies. First, genotypes for all animals typed at low density were imputed to higher density (N = 683,752 (ARS-UCD1.2) and 622,699 (UOA_Angus_1) SNP) using 1,166 reference animals with BovineHD-derived genotypes. In a second step, the partially imputed high-density genotypes were imputed to the sequence level using a reference panel of 161 sequenced animals. Both steps of imputation were carried out with Beagle 5.1 [72]. Variants with MAC *>* 3 (or) deviating significantly from Hardy-Weinberg proportions (P *<* 10^−6^), (or) with imputation accuracy (Beagle R^2^) less than 0.5 were filtered out. The imputed data with variants aligned to the ARS-UCD1.2 and UOA_Angus_1 assembly respectively, contained genotypes at 12,761,165 and 12,602,069 sequence variants.

### Genome-wide association testing and estimation of genomic heritability

We tested the association between phenotypes in the form of de-regressed proofs for six traits and sequence variants in between 11,294 and 12,434 BSW cattle. We considered phenotypes for stature (N=11,294), milk yield (N=13,388), protein yield (N=12,392), fat yield (N=12,388), protein content (N=12,439), and fat content (N=12,434). The SNP-based association study was carried out using a linear mixed model implemented with the MLMA-approach of the GCTA software package [73]. The model included a genomic relationship matrix built from 560,777 autosomal SNPs that were typed on the BovineHD chip (positions mapped according to ARS-UCD1.2) and four principal components to account for relatedness and population stratification. The genomic heritability was estimated for the six traits using the genomic restricted maximum likelihood (GREML) approach implemented in GCTA [73]. Therefore, we used genomic relationship matrices (GRM) that were built from all imputed autosomal sequence variants. We also partitioned the genomic heritability onto individual autosomes using GRM built from variants of the respective autosomes.

## Supplementary information

**Supplementary files**

Supplementary File 1: Table S1 — BSW cattle IDs Accession IDs of the 161 bovine samples used for our study.

Supplementary File 2: Table S2 — Number of mapped reads contained in the original files but not considered for our study.

Number of reads mapped to sexual chromosomes and to unplaced contigs for both assemblies. Low quality mapping includes the number of reads filtered out when considering only uniquely mapped properly paired reads with a mapping quality threshold of 10. Sample-wise mean and standard deviation can be found between parentheses. The length of the sexual chromosomes and unplaced contigs is also included.

Supplementary File 3: Table S3 — Number of variants during the different filtering steps: from original variants to high-quality and non-fixed variants.

Original variants are considered as the raw variants retrieved from GATK. Low quality variants are discarded during hard-filtering and fixed variants are identified when the minor allele count (MAC) is set to 1 in VCFtools. The percentage of variants to the original variants before hard-filtering are in parentheses.

Supplementary File 4: Fig. S1 — Variant density of the autosomes for both assemblies.

Number of variants detected per kilo base pair (Kb) along autosomal sequences of 161 BSW samples when aligned to the ARS-UCD1.2 (blue) and UOA_Angus_1 (orange) assembly.

Supplementary File 5: Fig. S2 — Density of variants across chromosomes 13 and 23.

The number of variants is shown within non-overlapping windows of 10 Kb for chromosome 13 (A) and 23 (B). The x-axis indicates the length of the chromosome (in Mb). The number of variants within each 10 Kb window is shown on the y-axis. Assembly ARS-UCD1.2 is displayed in the top panel (blue) and assembly UOA_Angus_1 is displayed as a mirror image in the bottom panel (orange).

Supplementary File 6: Fig. S4 — Density of high-quality and non-fixed variants per Kb along the autosomal genome.

Unlike Table 3 in the main text, densities are calculated here when chromosome 12 is not considered.

Supplementary File 7: Table S5 — Number and percentage of multiallelic variants.

Percentage of multiallelic variants is obtained from the division of multiallelic variants to non-fixed high-quality variants. Multiallelic variants are identified when the ‘–min-alleles 2 –max-alleles 2’ flag is set in VCFtools. Alleles not in Hardy-Weinberg proportions are the number of variants with P-value below the threshold of 10^−8^ when testing for Hardy-Weinberg proportions with PLINK. Percentages are between parentheses.

Supplementary File 8: Fig. S3 — Density of variants deviating from Hardy-Weinberg proportion for chromosome 12.

The number of variants differing from Hardy-Weinberg proportion are plotted as non-overlapping windows of 10 Kb along the autosomal sequence. The y-axis relates the variant density, number of variants per 100 Kb, for each 10-Kb-windows.

Supplementary File 9: Table S6 — Summary of the annotated sequence ontology classes of SNPs and INDELs.

SO terms are described by Ensembl. Total number of high-quality and non-fixed annotated SNPs and INDELs for both assemblies that were annotated using the release 101 annotation files with VEP tool.

Supplementary File 10: Table S7 — Candidate selection signatures detected using ARS-UCD1.2 as reference.

Genomic coordinates, CLR values, P-values and encompassed genes for 40 candidate selection signatures

Supplementary File 11: Table S8 — Candidate selection signatures detected using UOA_Angus_1 as reference.

Genomic coordinates, CLR values, P-values and encompassed genes for 33 candidate selection signatures

Supplementary File 12: Fig. S4 — Selective sweeps on chromosome 13.

Chromosome 13 region in ARS-UCD1.2 from 10,501,688 - 12,506,844 Mb and corresponding region on UOA_Angus_1 between 71,231,671 - 73,018,009 Mb with highlighted six selective sweep region from 11.5 Mb to 12 Mb. (A) Dot plot between the two assemblies, (B) SNP density per Kb (red line represents the average SNP density/chromosome), (C) Standardized coverage per 0.5 Kb, (D) Alternate allele frequency of each SNP (each dot is per SNP).

Supplementary File 13: Fig. S5 — Selective sweeps on chromosome 22.

Chromosome 22 region in ARS-UCD1.2 from 11,928,425 - 12,925,926 Mb and corresponding region on UOA_Angus_1 between 12,003,259 - 13,000,720 Mb with highlighted two selective sweep region. (A) Dot plot between the two assemblies, (B) SNP density per Kb (red line represents the average SNP density/chromosome), (C) Standardized coverage per 0.5 Kb, (D) Alternate allele frequency of each SNP (where each dot is per SNP).

Supplementary File 14: Fig. S6 — Genome Wide Association Study (GWAS).

Manhattan plots showing association of sequence variants — imputed using ARS-UCD1.2 (blue and grey) and UOA_Angus_1 (orange and grey) — with fat yield (A), protein yield (B) and stature (C).

## List of abbreviations

bp: Base pairs
BSW: Brown Swiss
CLR: Composite likelihood ratio
GATK: Genome Analysis Toolkit
GWAS: Genome-wide association study
h2: heritability
INDELs: insertions and deletions
Kb: Kilo base pairs
Mb: Mega base pairs
QTL: Quantitative trait locus
SNP: Single nucleotide polymorphism

## Ethics approval and consent to participate

Not applicable

## Consent for publication

Not applicable

## Availability of data and materials

Accession numbers for all animals are available in the Supplementary File 1: Table S1. All workflows and scripts used to produce the results are available at the ETH Animal Genomics Github repository [50].

## Competing interests

The authors declare that they have no competing interests.

## Funding

This study was supported by a grant from the Swiss National Science Foundation (310030 185229). The funding body was not involved in the design of the study and collection, analysis, and interpretation of data and in writing the manuscript.

## Authors’ contributions

Generated the data: HP, RF; Conceived and designed the experiments: ALV, HP; Analyzed the data: ALV, MB, NKK, HP; Wrote the manuscript: ALV, MB, NKK, HP. All authors have agreed on the content of the manuscript.

## Acknowledgements

We thank the Functional Genomics Center Zurich for generating DNA sequencing data. We thank the Arbeitsgemeinschaft Deutsches Braunvieh and the Arbeitsgemeinschaft Süddeutscher Rinderzüchter und Besamungsorganisationen e.V. (ASR) for granting access to whole-genome sequencing data of important key ancestor animals.

## Notes

https://github.com/AnimalGenomicsETH/Reference_assembly_choice

### Competing Interest Statement

The authors have declared no competing interest.

